# Dogs and humans share biomarkers of mortality

**DOI:** 10.1101/2025.08.20.671317

**Authors:** Benjamin R. Harrison, Joshua M. Akey, Noah Snyder-Mackler, Dan Raftery, DAP Consortium, Kate E. Creevy, Daniel E.L. Promislow

**Affiliations:** Department of Anesthesiology and Pain Medicine, University of Washington, Seattle, WA, USA; Lewis-Sigler Institute for Integrative Genomics, Princeton University, Princeton 08540, USA; School of Life Sciences, Arizona State University, Tempe, AZ, USA; Northwest Metabolomics Research Center, Department of Anesthesiology and Pain Medicine, University of Washington, Seattle, WA, USA; Department of Small Animal Clinical Sciences, Texas A&M University, College Station, TX, USA; Jean Mayer USDA Human Nutrition Research Center on Aging, Tufts University, Boston, MA, USA

## Abstract

There is growing interest in the use of molecular features as predictors of age, age-related disease risk and age-related mortality. A major shortcoming of this field, however, is the lack of suitable translational research models to identify and understand the underlying mechanisms of these predictive biomarkers in human populations. In particular, we lack a system which, like humans, is genetically variable, lives in diverse environments, and experiences aging-related chronic conditions treated in the context of a sophisticated health care system. Here, we present results from our analysis of data from the Dog Aging Project, a long-term longitudinal study of aging in more than 50,000 companion dogs. In particular, using longitudinal survival models, we present the striking finding of a strong, highly significant positive correlation between the effect of individual metabolites on all-cause mortality in humans, and the association of those same metabolites on all-cause mortality in dogs. We also find that across these independent human studies, the biomarkers identified are also highly correlated, strongly suggesting a general signature of mortality within the plasma metabolome across humans, and now in dogs as well. Given the many similarities between dogs and humans with respect to genetics, environment, disease, and disease treatment, and the fact that dogs are so much shorter lived than humans, we argue that dogs represent an extremely valuable translational model in our ongoing effort to understand the underlying molecular causes and consequences of age-related morbidity and mortality in humans.

## Introduction

In an effort to predict and explain variation in morbidity and mortality, researchers have turned to the metabolome, which consists of the small molecules that make up the structural and functional building blocks of life. Numerous human cohort studies have now shown that concentrations of specific metabolites in the blood are associated with future risk of mortality (1–5). However, these studies are enormously challenging. A sample of thousands of individuals must be followed for a decade or more, at great expense, to determine who dies and at what age. As an alternative, companion dogs exhibit considerable genetic variation, share human-built environments, and suffer from many of the same age-related conditions as humans (6), yet are much shorter lived.

They thus offer a powerful translational model of aging. Here we mapped associations between all-cause mortality and the blood metabolome in a focused cohort, the Precision cohort (7), within a longitudinal study of companion dogs, the Dog Aging Project (8,9).

The Precision Cohort of nearly one thousand companion dogs across the United States includes a wide range of dog breeds, with over half of dogs being of mixed breed ancestry (7). The design of the Precision Cohort is such that their sex and sterilization status, the distribution of urban, suburban and rural homes, and broader geographic representation reflects that of companion dogs in the US. We have data from both owner surveys and annual clinical visits for the Precision Cohort which enable longitudinal analysis of mortality and other health outcomes (7,8).

In the first longitudinal analysis of mortality within the Precision Cohort, we find that biomarkers of mortality in the blood metabolome of dogs are highly concordant with those across a breadth of human cohorts, and have been identified in a much shorter timeframe. This clearly demonstrates the potential of companion dogs as a translational aging model.

## Results

Using plasma samples collected from 937 dogs enrolled in the Precision Cohort, we used a longitudinal model to determine the ability of 133 metabolites to predict all-cause mortality while controlling for age, sex, weight, and breed/genetic relatedness. From the distribution of P-values, we estimated that ∼17% of metabolites are associated with mortality, of which 23 are significant at an FDR < 5% (Figure 1a, see Methods).

**Figure 1.**
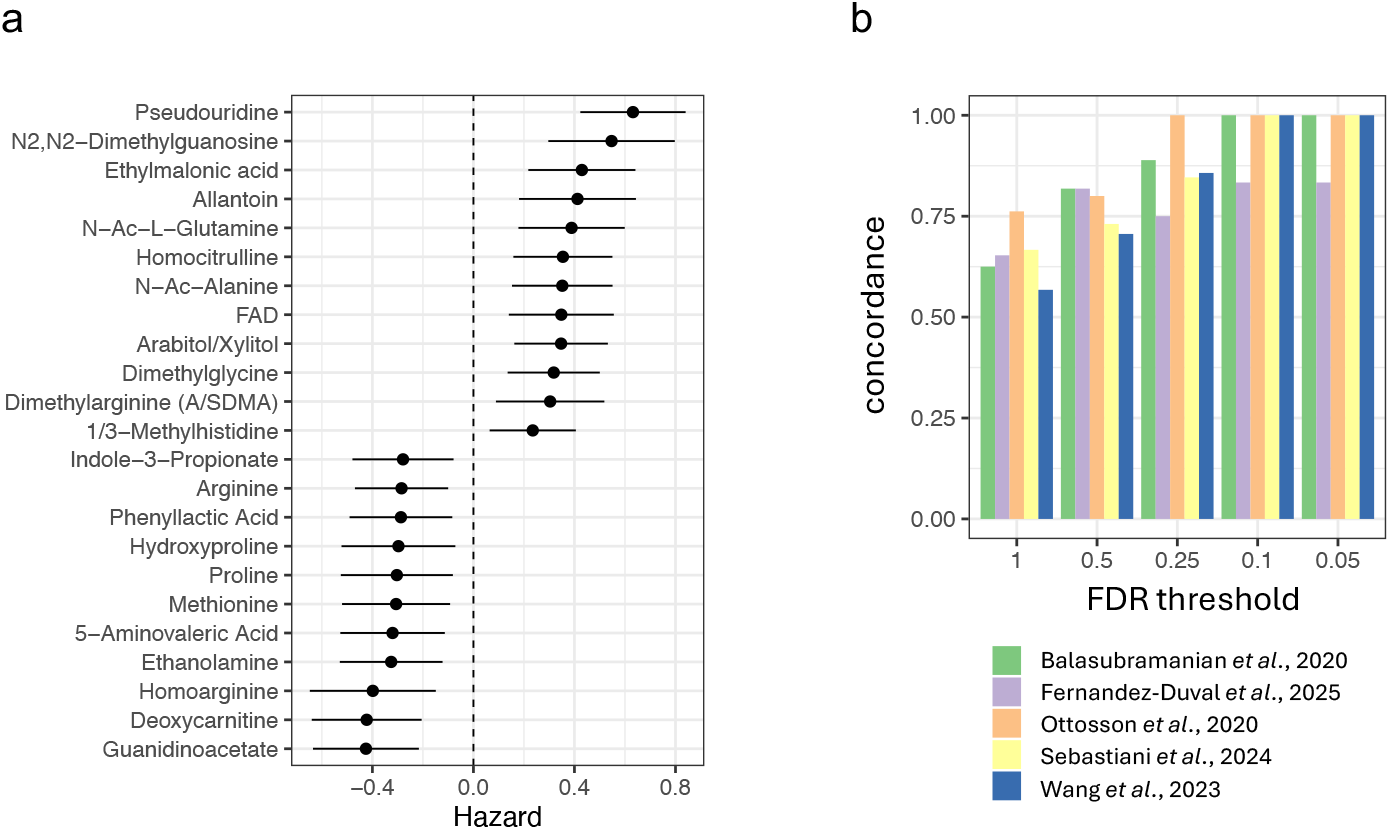
Similarity of the mortality-associated plasma metabolome in dogs and humans. (**a**) All-cause mortality hazard, with 95% confidence intervals, for 23 metabolites in dog plasma (FDR ≤ 0.05). Hazard>0 (<0) indicates greater (lesser) mortality risk. (**b**) The concordance of hazard ratio (HR) between dogs and humans for 37 to 64 plasma metabolites measured in each of five human studies over several FDR thresholds. The null expectation of concordance is 0.5.

We then tested whether metabolites predictive of mortality in dogs were also associated with mortality in human studies of plasma biomarkers (1–5), first comparing the concordance of the effects, and then looking at the overall correlation across hazard ratios (HR) (see Methods). In a set of five independent human studies, we found between 37 and 64 metabolites that were also measured in the dogs, and the sign of the HR in humans and dogs was concordant in 64% of these instances (Fisher’s exact test P = 9.3 × 10^-9^). The concordance in HR among metabolites measured in dogs and in each of five human studies rose substantially when we limited the analysis to metabolites most strongly associated with mortality, reaching perfect concordance for most studies at an FDR <0.1 (Figure 1b, and Supplementary Information).

This level of concordance suggested that not only might dogs share biomarkers of mortality with humans, but that across these five independent human studies, a shared signature of mortality risk could be identified. We identified between 24 and 45 (mean=31.8) metabolites measured in any two human studies. Among all pairs of studies, the association of intersecting metabolites with mortality were correlated from at least r=0.37 to 0.85 (with P<0.05 in all cases, Figure 2a). We also found HR in humans strongly correlated with HR in dogs, with Pearson’s r ranging from 0.46 to 0.74 (P < 0.002 in all cases; Fisher’s combined probability of 4.0 x 10^-17^, Figure 2a, Supplementary Information).

**Figure 2.**
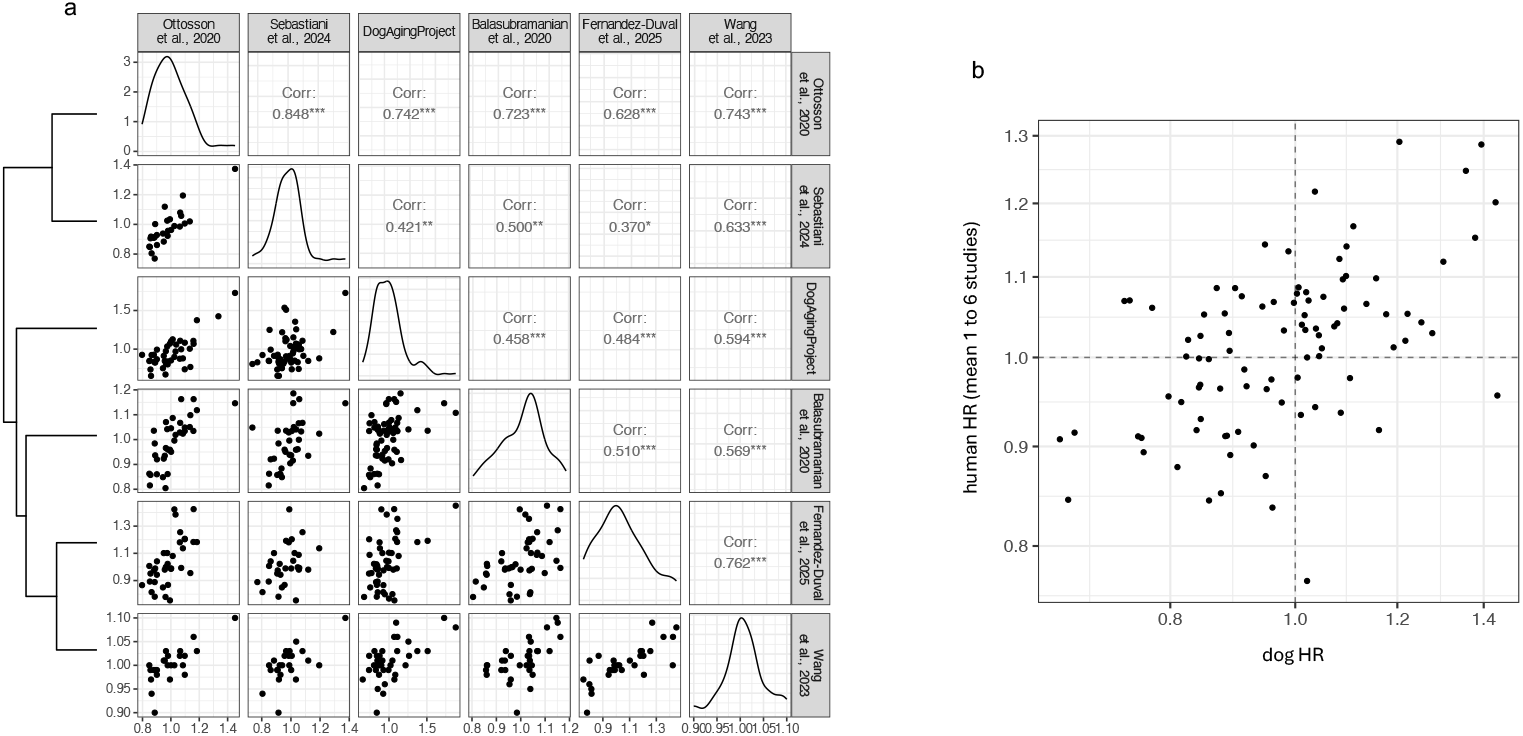
Pairwise analysis of mortality associated with metabolites identifies mortality biomarkers shared generally among humans and dogs. (**a**) The lower triangle shows pairwise plots of the HR associated with each of 37 to 64 metabolites measured in the Precision Cohort of the DAP, and in each of five human studies. The diagonal shows the distribution of HR within each study. Above the diagonal are pairwise Pearson’s correlation (^*^P<0.05, ^**^P<0.005, ^***^P<0.0005). The dendrogram on the left shows the clustered pairwise similarity in HRs among the studies. (**b**) The geometric mean of up to six HRs measured per metabolite (from five studies in **a** and four additional human studies without sufficient intersecting metabolites to test them individually for concordance), versus the HRs in dogs (Pearson’s r = 0.52, P = 8.9x10^-8^).

There were four additional human studies that reported HR for ten or fewer of the metabolites that were measured in the dogs and so were underpowered for these comparisons. To include those studies, we aggregated data for each metabolite from all nine human studies, by taking the geometric mean HR per metabolite. There were mean HRs for 93 metabolites that were measured in an average of 2.9 human studies (range 1 to 6) and in the dogs. These mean human HRs correlate highly with the HRs in dogs (r = 0.52, P = 8.9 × 10^-8^, Figure 2b).

Among the mortality associations seen in both dogs and humans, we found higher risk associated with more abundant pseudouridine, N2,N2-dimethylguanosine and homocitrulline, and with less abundant deoxycarnitine and homoarginine (Figure 1a). The abundance of these metabolites in blood associates with renal function in humans (e.g., 10), suggesting a potential shared physiological axis of aging in dogs and humans.

The mortality analysis in the Precision Cohort involved 104 deaths among 937 dogs that were followed for an average of 2.6 years (maximum = 3.9 years). By contrast, average follow-up times among participants reported in the human studies ranged from 8.0 to 22.5 years and included cohorts that have been followed for as long as 28 years (Supplementary Information). Thus, metabolites associated with all-cause mortality in dogs were not only significantly correlated with the same metabolites associated with all-cause mortality in humans but were also identified substantially faster.

## Discussion

Here we have identified strong metabolite predictors of all-cause mortality and demonstrated the potential of companion dogs as a translational aging model. Studies of human mortality and the plasma metabolome have been conducted using thousands of participants. The studies we incorporated into our analysis involve over 24 cohorts, with representation from both sexes, several ethnic groups, and both healthy and unhealthy individuals (Table S1). These human studies independently survey for plasma metabolites associated with mortality, and across studies we find these biomarkers are substantially correlated. This result indicates that humans, generally, share a signature of mortality risk within the blood metabolome. We also demonstrated that all-cause mortality in companion dogs is associated with a very similar profile of metabolites as is in humans. Importantly, the speed with which we were able to identify biomarkers of mortality in dogs underscores the high translational potential of the dog as a model of aging.

The blood metabolites associated with mortality in humans typically associate with glomerular filtration rate (10,11), which is consistent with the filtration and resorption of metabolites at the blood and urine interface (12). Here we show that some of the same metabolites also associate with mortality in dogs. These results suggest that the decline in kidney function with age in humans may explain the association of these metabolites with mortality. We controlled for age, weight and serum creatinine among the dogs to control for the effects of size on kidney function and the portion of glomerular filtration reflected in creatinine (13).

As we continue to follow dogs enrolled in the Dog Aging Project through all life stages, we stand to identify biomarkers from additional -omic domains, with the power to predict a wide range of age-related diseases as well as mortality. This work can help us to understand the mechanistic underpinnings of variation in aging in both dogs and in us, and to realize the potential of geroscience to identify healthspan interventions within our lifetimes (14).

## Materials and Methods

The Dog Aging Project is a long-term longitudinal study of the genetic and environmental determinants of healthy aging in more than 50,000 US-based companion dogs. Within this study population, a subset of 937 dogs, the ‘Precision Cohort’, have contributed biospecimens with annual follow-ups from which we obtain complete blood count and blood chemistry, urinalysis, fecal microbiome, blood cell epigenome, flow cytometric analysis of white blood cells, and the focus of this current study, the plasma metabolome (7).

Dog sex, weight and live/dead status were obtained from annual owner survey data on the Terra server (https://terra.bio). Metabolome data were obtained by targeted liquid chromatography-tandem mass spectrometry (LC-MS/MS) of aqueous metabolites extracted from plasma samples (7). Peak-areas for 133 metabolites were log-normalized, mean-centered, adjusted for effects of extraction batch, LC-MS run order, sample hemolysis, sample lipemia, sample arrival temperature, and fasting-time prior to blood collection. Missing data were imputed using a custom elastic net procedure (https://github.com/ben6uw/Dog-Aging-Project-Publication-Resources/tree/main/dog_human_mortality).

For mortality analysis, we used a time-dependent mixed-effects Cox proportional hazards model to analyze time-to-death (from any cause), where ‘time 0” was the date of the first veterinary visit where blood was collected. Observations were censored at the last date of contact (either survey or blood collection date) if the dog was still alive. We adjusted for age at ‘time 0’, the square-root of weight at ‘time 0’, sex, and scaled log_e_-serum creatinine as fixed effects, with scaled plasma metabolites and scaled log_e_-serum creatinine as time-varying covariates that were updated at each annual follow-up blood collection. We also included a random effect of a genome-relatedness matrix (https://terra.bio). We accounted for multiple testing by controlling the false discovery rate(15).

The nine human studies we included were identified in literature searches. We identified several additional studies of human mortality but excluded them because they appeared to share the same metabolome data as one of the studies we included. The human mortality associations were obtained from one or more Cox models in each of the nine studies. For eight of the human studies, results for Cox models were adjusted for age and other covariates including sex, body mass index, smoking status, hypertension, medications for hypertension and hyperlipidemia, diabetes status, heart disease, cholesterol, as well as sub-cohorts and case/control status were used when applicable. When available, Cox models with full adjustments were used to compare HR among human studies with dog HR, although using human HRs from simpler models did not substantially affect the results.

Data and R code used to analyze these data and create figures are all available at our GitHub site: https://github.com/ben6uw/Dog-Aging-Project-Publication-Resources/tree/main/dog_human_mortality.

## Supporting information

human cohort summary

## Acknowledgments and Funding

We thank Mandy Kauffman, Maria Partida-Aguilar, Paul Litwin, Fillipo Artoni, Tom Murdy and Jessica Foley for helpful comments on an earlier draft of this manuscript. This research is based on publicly available data collected by the Dog Aging Project, under U19 grant AG057377 (PI: Daniel Promislow) from the National Institute on Aging, a part of the National Institutes of Health, and by additional grants and private donations, including generous support from the Glenn Foundation for Medical Research, the Tiny Foundation Fund at Myriad Canada, the WoodNext Foundation, and the Dog Aging Institute. DP received support from USDA cooperative agreement USDA/ARS 58-8050-9-004.

## Supplementary Information

**Supplementary Figure 1.** Reproducible concordance in the mortality associated plasma metabolome in dogs and humans

**Supplementary Table 1.** Cohort Summary

