## Supplemental Material for "Dogs and humans share biomarkers of mortality"

**Supplemental Information**

**Supplemental Figure 1**

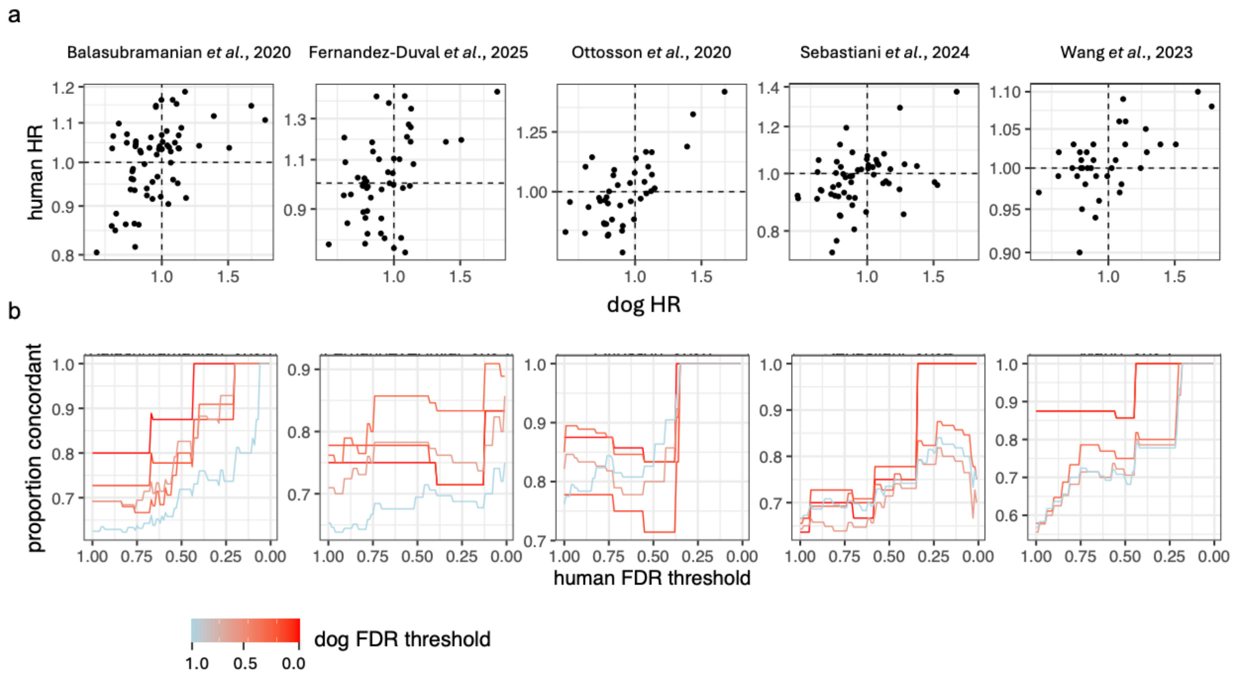


**Supplementary Figure 1. Reproducible concordance in the mortality associated plasma metabolome in dogs and humans.** (**a**) The hazard ratio (HR) of 37 to 64 plasma metabolites in five human studies (top row), versus the hazard ratios of the same plasma metabolites in the Precision Cohort of the Dog Aging Project (dog HR). (**b**) Metabolites with more significant mortality associations exhibit higher concordance in dog-human comparisons. Among each of five subsets of dog metabolites, limited by increasingly significant associations with dog mortality (dog FDR threshold, color scale), the proportion of metabolites whose HR is concordant with humans (both HRs > 1 or both HRs < 1) rises with the stringency of association in humans (human FDR threshold).

**Supplementary Table 1. Cohort Summary**

[attached PDF]
