## Supplementary material for "Dogs and humans share biomarkers of mortality": human cohort summary

| PMID | study | cohort | sex (%female) | age | N subjects | deaths | freq. death | follow up (yrs) | follow up measure |
|---|---|---|---|---|---|---|---|---|---|
| this study | this study | Precision Cohort of the Dog Aging Project | 48.6 | mean=5.3 (SD=3.3) | 937 | 104 | 0.11 | 2.64 | average |
| 31431621 | Deelen et al 2019 | Pravastatin in elderly individuals at risk of vascular disease | 0.515 | 75.3 | 5329 | 467 | 0.09 | 2.76 | average |
| 31431621 | Deelen et al 2019 | Leiden Longevity Study | 0.613 | 97.35 | 843 | 823 | 0.98 | 4.03 | average |
| 31431621 | Deelen et al 2019 | United Kingdom Adult Twin Registry | unknown | 64.58 | 1996 | 58 | 0.03 | 4.32 | average |
| 31431621 | Deelen et al 2019 | Avon Longitudinal Study of Parents and Children | unknown | 47 | 4351 | 17 | 0.00 | 5.69 | average |
| 32751974 | Ottosson et al 2020 | Malmö Preventive Project | 23.6 | 70.5 | 369 | 118 | 0.32 | 7.7 | average |
| 31431621 | Deelen et al 2019 | Dietary, Lifestyle, and Genetic determinants of Obesity and Metabolic syndrome | 0.532 | 52.39 | 4816 | 190 | 0.04 | 7.73 | average |
| 31431621 | Deelen et al 2019 | Alpha Omega Cohort | 0.246 | 69.21 | 568 | 157 | 0.28 | 7.79 | average |
| 31431621 | Deelen et al 2019 | Estonian Biobank | 0.626 | 46.1 | 10,988 | 912 | 0.08 | 7.97 | average |
| 31431621 | Deelen et al 2019 | Cooperative Health Research in the Region of Augsburg | 0.513 | 60.89 | 1790 | 123 | 0.07 | 8.02 | average |
| 31431621 | Deelen et al 2019 | The Rotterdam Study | 0.581 | 75 | 2963 | 1254 | 0.42 | 8.28 | average |
| 31651959 | Balasubramanian et al 2020 | Women's Health Initiative Hormone Therapy | 100 | range 50 to 180 | 1355 | 685 | 0.51 | 9.1 | median |
| 31651959 | Balasubramanian et al 2020 | Women's Health Initiative Observational Study | 100 | range 50 to 180 | 943 | 417 | 0.44 | 10.6 | median |
| 31431621 | Deelen et al 2019 | Erasmus Rucphen Family Study | 0.549 | 50.44 | 680 | 107 | 0.16 | 10.67 | average |
| 31431621 | Deelen et al 2019 | Leiden Longevity Study | 0.554 | 70.93 | 2241 | 191 | 0.09 | 11.76 | average |
| 40107652 | Fernández-Duval et al 2025 | Mediterranean diet for primary prevention of cardiovascular diseases | 57.5 | range 55 to 80 | 1878 | 457 | 0.24 | 12.2 | median |
| 39504246 | Sebastiani_2024 | Long Life Family Study | 54 | 24 to 110, median 74 | 1267 | unknown | NA | 15 | minimum |
| 31431621 | Deelen et al 2019 | National FINRISK Study | 0.503 | 48.29 | 7603 | 1213 | 0.16 | 16.7 | average |
| 25864806 | Cheng et al 2015 | Framingham Offspring Study | unknown | unknown | 2,327 | 439 | 0.19 | 17.4 | total |
| 32751974 | Ottosson et al 2020 | Malmö Diet and Cancer-Cardiovascular Cohort | 44.9 | 59.5 | 374 | 180 | 0.48 | 18.3 | average |
| 26956554 | Yu et al 2016 | Atherosclerosis Risk in Communities | 57.7 | ~52-56 (range of means) | 1887 | 671 | 0.36 | 22.5 | average |
| 37717037 | Wang et al 2023 | Health Professional Follow-Up Study | 0 | unknown | 1,620 | 4288* | NA | 22.60 | median |
| 37717037 | Wang et al 2023 | Nurses' Health Study I | 100 | unknown | 6,883 | 4288* | NA | 22.60 | median |
| 37717037 | Wang et al 2023 | Nurses' Health Study II | 100 | unknown | 3,131 | 4288* | NA | 22.60 | median |
| 29390044 | Huang et al 2018 | Alpha-Tocopherol, Beta-Carotene Cancer Prevention | 0 | range 50 to 69 | 620 | 435 | 0.70 | 28 | total |
